# How Competition Affects Evolutionary Rescue

**DOI:** 10.1101/022202

**Authors:** Matthew Miles Osmond, Claire de Mazancourt

## Abstract

Populations facing novel environments can persist by adapting. In nature, the ability to adapt and persist will depend on interactions between coexisting individuals. Here we use an adaptive dynamic model to assess how the potential for evolutionary rescue is affected by intra- and interspecific competition. Intraspecific competition (negative density-dependence) lowers abundance, which decreases the supply rate of beneficial mutations, hindering evolutionary rescue. On the other hand, interspecific competition can aid evolutionary rescue when it speeds adaptation by increasing the strength of selection. Our results clarify this point and give an additional requirement: competition must increase selection pressure enough to overcome the negative effect of reduced abundance. We therefore expect evolutionary rescue to be most likely in communities which facilitate rapid niche displacement. Our model, which aligns to previous quantitative and population genetic models in the absence of competition, provides a first analysis of when competitors should help or hinder evolutionary rescue.

## 1 Introduction

Individuals are often adapted to their current environment [1]. When the environment changes individuals may become maladapted, fitness may drop, and population abundances may decline [2]. If the changes in the environment are severe enough, populations may go extinct. But populations can also evolve in response to the stress and thereby return to healthy abundances [3, 4]. Why some populations are capable of rescuing themselves from extinction through evolution, while others go extinct, is a central question to both basic evolutionary theory and conservation [5].

Ecological and evolutionary responses to changing environments are contingent on the community in which the change occurs [6, 7, 8, 9, 10]. A population’s ability to adapt and persist in changing environments will therefore also hinge on the surrounding community [11] (see also [12], this issue). By including the ecological community in a formal theory of adaptation to changing environments, we may better predict the response of natural communities to contemporary stresses, such as invasive species [13, 14] and global climate change [15, 16].

Competition reduces population abundance [17, 18, 19, 20]. Since less abundant populations are more likely to go extinct when exposed to new environments [21, 22], competition may therefore lower the potential for evolutionary rescue. But competition can also increase selective pressure [23], speed niche expansion [24, 25, 26], and increase rates of evolution [27], possibly allowing populations to adapt to new conditions faster. These potentially contrasting effects may account for the unanticipated population dynamics and patterns of persistence in competitive communities [6] (but see [10]).

Currently, most theory on adaptation to abrupt environmental change consider only isolated populations [3, 28, 29, 30, 31, 32, 33], and many of these studies assume unbounded population growth, thus ignoring intraspecific competition as well. The studies that do consider intraspecific competition, in the form of negative density-dependence, give inconsistent conclusions, stating that density-dependence has no effect [29] or decreases [30, 34] persistence. Of the handful of studies that examine the effect of interspecific competition on adaptation to environmental change, nearly all predict slower adaptation and more extinctions (reviewed in [35]). One notable exception suggests that interspecific competition can aid persistence in a continuously changing environment, by adding a selection pressure that effectively “pushes” the more adapted populations in the direction of the moving environment [36].

Here we use the mathematical framework of adaptive dynamics to describe the evolutionary and demographic dynamics of a population experiencing competition and an abrupt change in the environment. Adaptive dynamics allows us to incorporate both intra- and interspecific competition in an evolutionary model while maintaining analytical tractability. We assess the potential for evolution to rescue populations by measuring the “time at risk”, i.e. the time a population spends below a critical abundance [3]. First, we derive an expression for the “time at risk” in a population undergoing an abrupt change in isolation. We then compare our results to previous studies and test the robustness of our results by relaxing a number of simplifying assumptions using computer simulations. Finally, we examine how a population’s ability to adapt and persist to an abrupt environmental change is impacted by the presence of competing species.

## 2 Model and Results

### 2.1 One-population model

We first examine how, in the absence of competitors, an asexual population with density- and frequency-dependent population growth responds to an abrupt change in the environment.

We assume that each individual in the population has a trait value *z*, and that a phenotype’s growth rate is determined by both its own trait value as well as the trait value of all other individuals within the population. Population dynamics are described by the logistic equation (Equation 2 in [37])

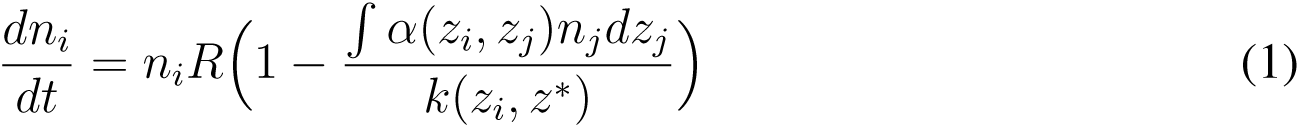

where *n*_*i*_ is the number of individuals with trait value *z*_*i*_, *R* is the per capita intrinsic growth rate, *a*(*z*_*i*_*, z*_*j*_) is the per capita competitive effect of individuals with trait *z*_*j*_ on individuals with trait *z*_*i*_, and *k*(*z*_*i*_*, z*^*\**^) is the carrying capacity of individuals with trait *z*_*i*_ in an environment where the trait value giving maximum carrying capacity is *z*^*\**^. We describe carrying capacity *k* as a Gaussian distribution (Equation 1 in [37])

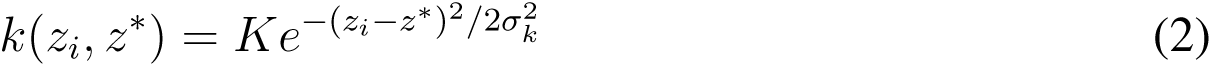

where *K* is the maximum carrying capacity and *σ*_*k*_ > 0 is the ‘environmental tolerance’, which describes how strongly carrying capacity varies with *z*_*i*_. For a given deviation from *z*^*\**^, smaller variances 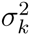 mean larger declines in carrying capacity *k*. We therefore refer to 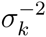 as the strength of stabilizing selection. Data on yeast responses to salt [5, 38] fit Gaussian carrying capacity functions, as described by Equation 2 (ESM).

We do not give a specific form for intraspecific competition *α*, but instead give requirements that are satisfied by a wide range of functions. First, we assume that individuals with the same trait value compete most strongly, that is 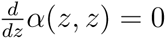 and 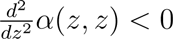. This is biologically reasonable and could describe, for instance, the effect of beak size on finches competing for seeds, where individuals with similar sized beaks compete strongly for similar sized seeds [39]. And we abitrarily set *α*(*z, z*) = 1, meaning that individuals with the same trait value take up one “unit” of carrying capacity.

Trait value *z* is assumed to be determined by a large number of loci, each with equal and small effect, making the range of possible phenotypes continuous and unbounded (i.e., *z* ∊ ℝ). To proceed analytically, we first assume that mutations are rare. The population remains monomorphic, with all individuals having “resident” trait value ẑ. The evolutionary trajectory is determined by the per capita growth rate of rare mutants in the neighborhood of ẑ (adaptive dynamics; [40]). When mutations are sufficiently rare, evolution occurs slow enough for us to consider the population at demographic equilibrium on an evolutionary timescale. This stands in contrast to previous models which jointly model demography and evolution (e.g., [3, 34]). The timescale separation between demography and evolution allows us to incorporate intra- and interspecific competition while maintaining analytical tractability. We later use computer simulations to examine how our analytical results perform when demography and evolution occur on similar timescales.

In Appendix A we show that when 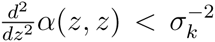 the ‘optimal trait value’ *z*^*\**^ is both convergence stable (i.e., by small steps the resident trait converges to *z*^*\**^) and evolutionary stable (i.e., once 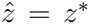 no other strategies can invade; *z*^*\**^ is an ESS, *sensu* Maynard Smith and Price [41]). We assume 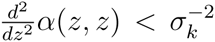 for the remainder of the paper, which means frequency-dependence is weak enough [42]. Our results apply for any function *α*, as long as *z*^*\**^ is both convergence and evolutionary stable.

Let our population begin in a constant environment with optimal trait value 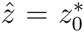. In time, all individuals become perfectly adapted 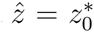. The population will reach equilibrium abundance *ñ* = *K* and its growth rate will become zero (Figure 1). Let us call this original abundance *K*_0_.

**Figure 1:**
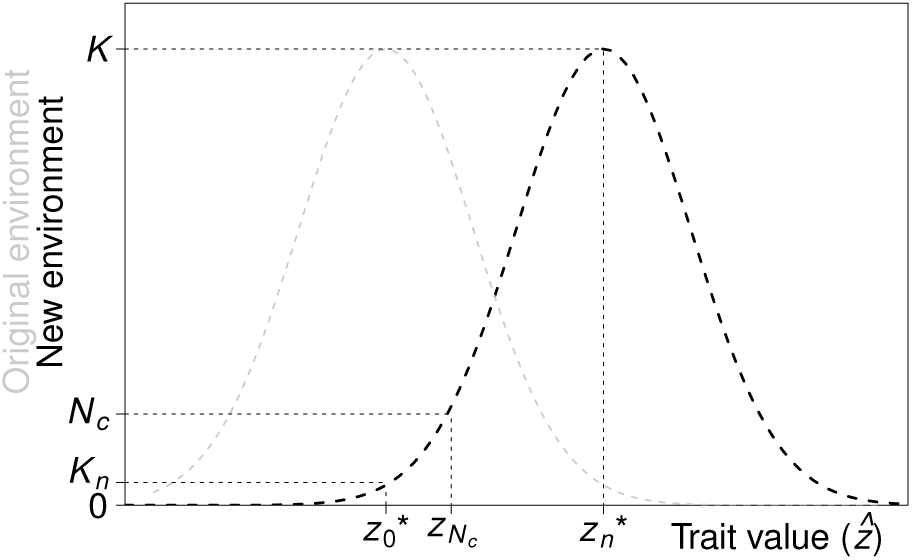
Our initially adapted population is monomorphic for the optimal phenotype in the original environment 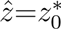(gray). When the environment changes, the carrying capacity function shifts (black). The new carrying capacity of our population 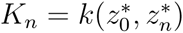 is the height of the intersection of the original trait value 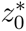 and the new carrying capacity function. The population evolves towards the new optimal phenotype 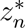 The population is at risk of extinction while its abundance is less than *N*_*c*_, or equivalently, while 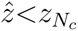

Suppose then that the environment suddenly changes so that the new optimal trait value is 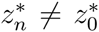. Our monomorphic population, with trait value 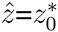, then immediately has equilib-rial abundance 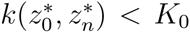 (Figure 1). The environmental change serves to decrease the carrying capacity of the population. The population will initially survive the abrupt change if 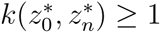 or, equivalently

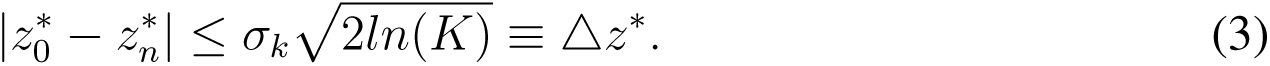

Note that setting *ñ* ≥ 1 as the extinction threshold scales population abundance in units of minimal viable population size [43, 37]. Because 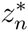 is the new evolutionary and convergence stable strategy, if the population survives the change it will evolve toward the new optimal trait value,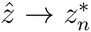. According to the canonical Equation of adaptive dynamics [44], the monomorphic trait value 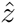 will change at rate

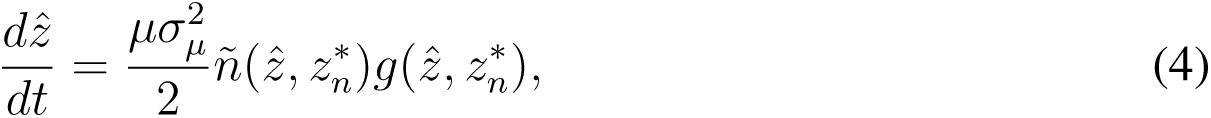

where *µ* is the per capita per generation mutation rate, 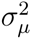 is the mutational variance (mutations symmetrically distributed with mean of parental value), 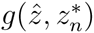 and is the local fitness gradient (Appendix A):

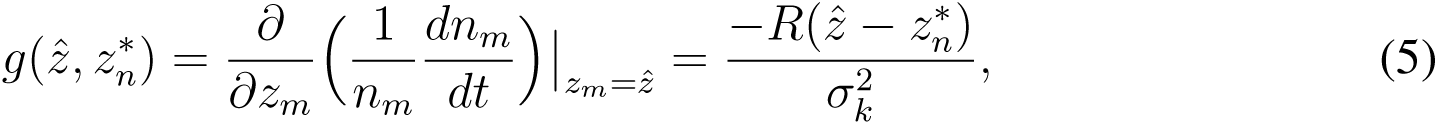

where *n*_*m*_ and *z*_*m*_ are a rare mutant’s abundance and trait value, respectively, and 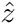 is the resident trait value [40]. The local fitness gradient describes the slope of the fitness function in the neighborhood of the parental trait value. Steeper slopes signify greater fitness differences between individuals with similar but unequal trait values [45]. Notice that 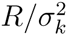 is the strength of stabilizing selection per unit time.

The rate of change in trait value is then:

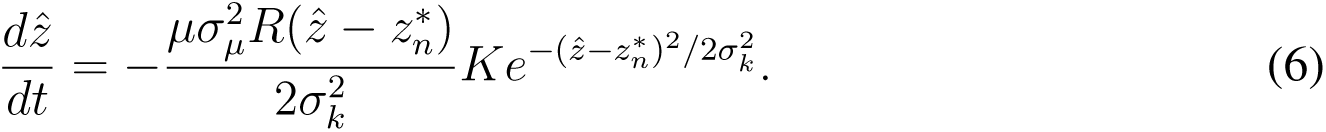

We cannot solve Equation 6 explicitly for 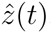 but using a first-order Taylor expansion we derive an approximate solution, describing evolution and demography following the abrupt change (Appendix B):

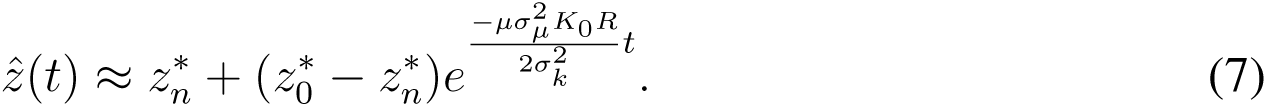

and

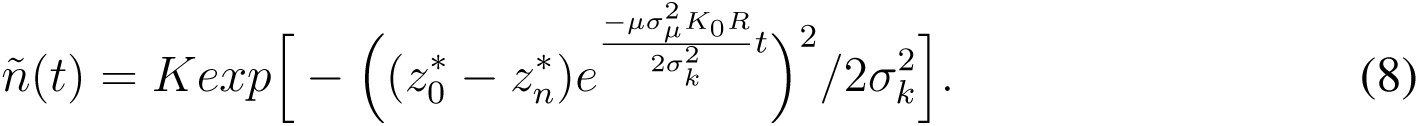

Taking the Taylor expansion about 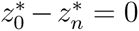 results in the assumption that the environmental change 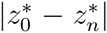 is small relative to environmental tolerance *σ*_*k*_ (i.e., a weak “initial stress”). Our first-order approximation of the Gaussian *k* is therefore taken at the maximum *z* = 0, which is a line with slope zero and height *K*_0_. This means we assume mutational input *µk* is constant at *µK*_0_, effectively decoupling the demographic and evolutionary dynamics of the recovering population. Our first-order approximation is the highest-order for which we can obtain an analytical solution.

Now, let *N*_*c*_ be the abundance below which demographic or environmental stochasticity are likely to cause rapid extinction [46, 3]. We use this heuristic *N*_*c*_, in the place of stochastic models, for simplicity. We are interested in the amount of time a population spends below this threshold, i.e., how long the population is at risk of extinction.

The population will never be at risk of extinction if its equilibrial abundance *ñ* remains above the critical abundance *N*_*c*_. In this model equilibrial abundance strictly increases in evolutionary time in a constant environment. Abundance is therefore at a minimum immediately following the abrupt shift in the environment. The population will avoid all chance of extinction if 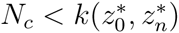 or, rearranging,

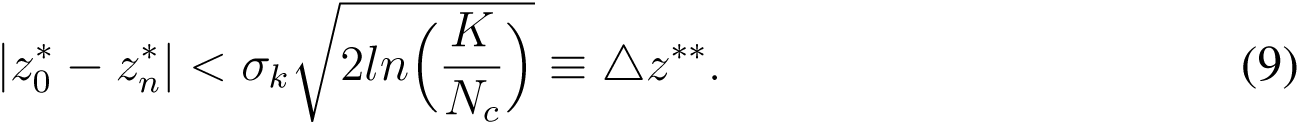

Here, we are most interested in the case where the population initially survives the abrupt change but abundance drops below the critical abundance 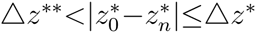, as this is when evolution is required to rescue populations from extinction.

−From Equation 2 we can find the trait value *z*_*N*_*c* required for a carrying capacity of *N*_*c*_. Plugging *z*_*N*_*c* into Equation 7 and solving for *t* gives the time it will take a population to evolve to this safe trait value *z*_*N*_*c*, which we will call the “time at risk” *t*_*r*_ (Figure 2)

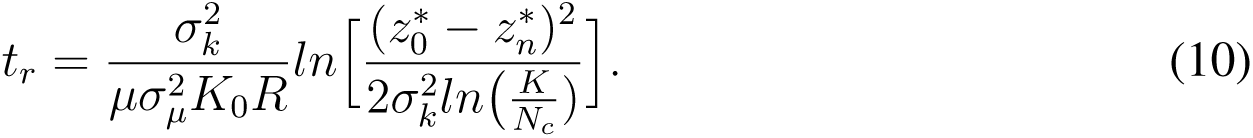

**Figure 2:**
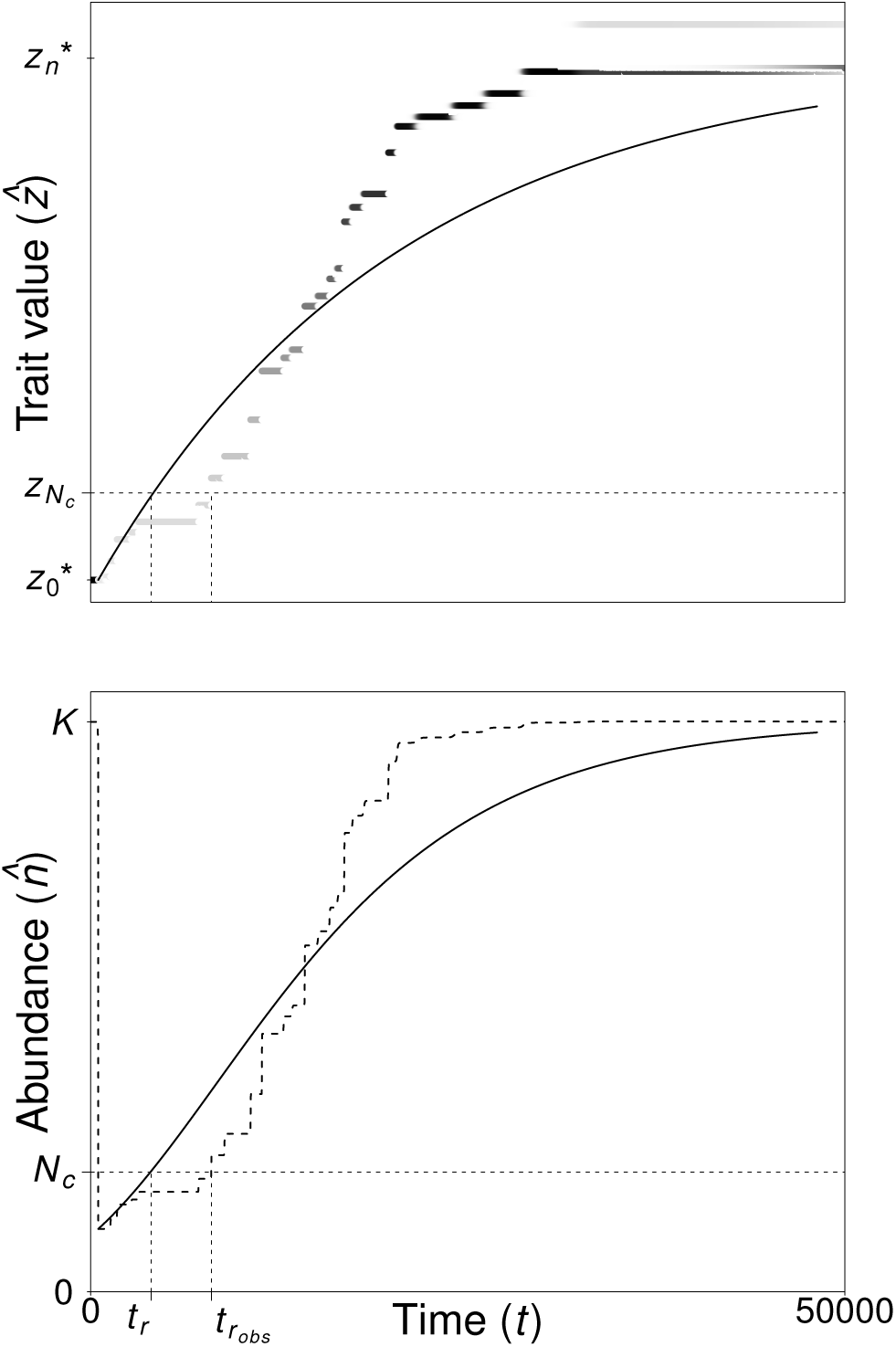
Adaptation following an abrupt change in the environment. (*Top*) Population trait value 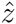 evolves towards the new optimal 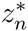(Equation 7). The time it takes to evolve a trait value *z*_*N*_*c*, which gives a critical abundance *N*_*c*_, is the expected “time at risk” *t*_*r*_ (Equation 10). (*Bottom*) Population abundance *ñ* increases as the population adapts to the new environment (Equation 8). Solid lines are analytical predictions (Equations 7 and 8). Greyscale is trait value weighted by abundance in a computer simulation, with dark common and white rare. The thick dashed line is total abundance at each time step in simulation. The observed time at risk is denoted *t*_*r*_*obs*.

So the time at risk *t*_*r*_ increases with the strength of the initial stress 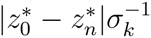 and the ratio of critical abundance to maximum carrying capacity *N*_*c*_*/K* and decreases with the mutational input *µK*_0_, mutational variance 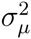, and the strength of stabilizing selection per unit time, 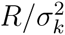 Time at risk *t*_*r*_ is a unimodal function of environmental tolerance *σ*_*k*_, with longest times at intermediate tolerances (Figure 3). Time at risk is reduced at small and large environmental tolerances because small tolerances cause strong selection (and hence fast evolution) and large tolerances allow greater abundances for a given degree of maladaptation.

**Figure 3:**
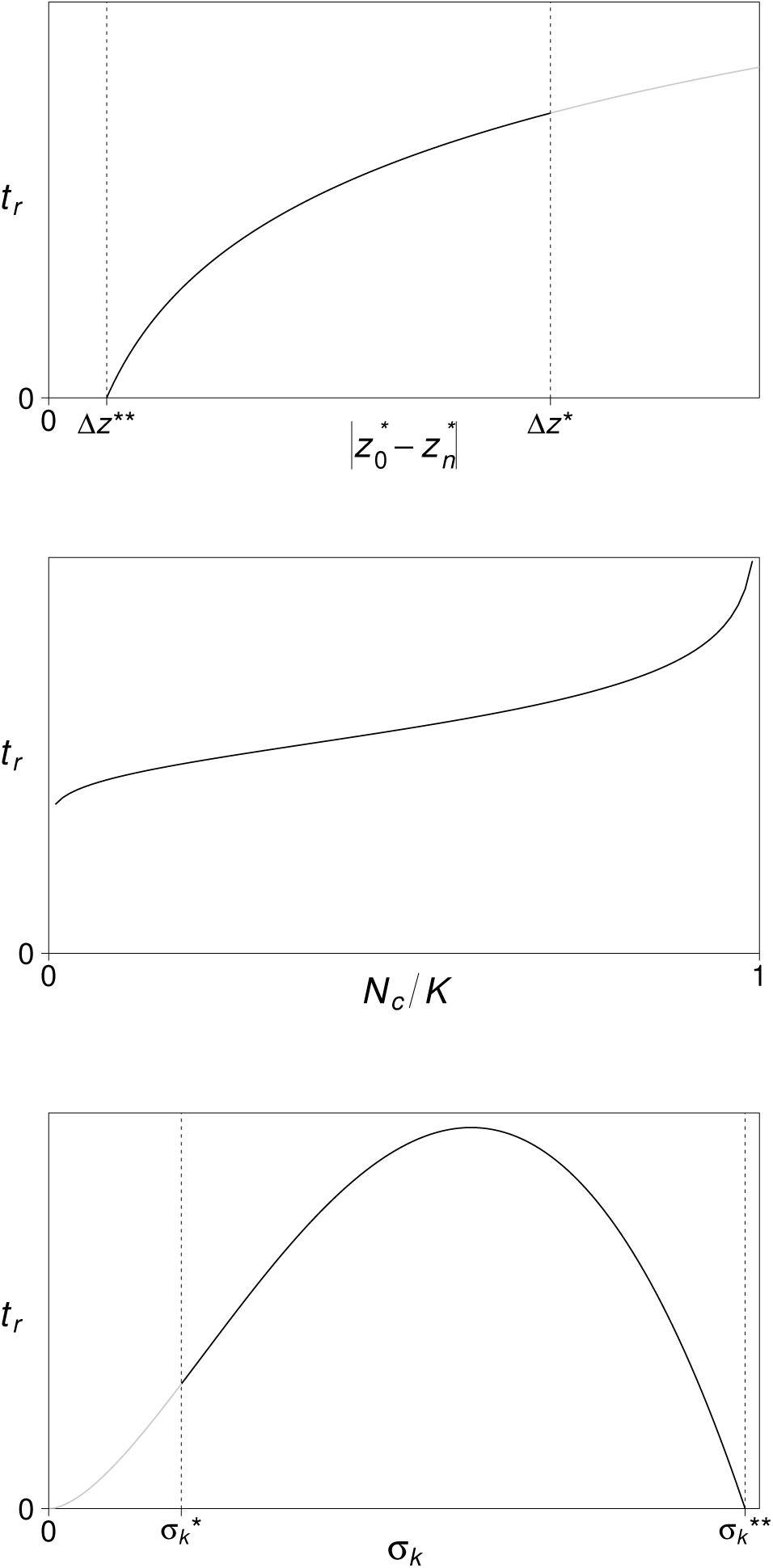
(*Top*) Time at risk *t*_*r*_ (Equation 10) increases monotonically with the magnitude of environmental change 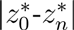 Magnitudes of change smaller than *Dz*^*\*\**^ are not large enough to put the population at risk of extinction (Equation 9) and magnitudes of change larger than *Dz*^*\**^ cause immediate extinction (Equation 3). (*Middle*) Time at risk *t*_*r*_ increases as the critical abundance *N*_*c*_ approaches maximum abundance *K*. As the critical abundance approaches the maximum abundance, *N*_*c*_*/K* → 1, the ratio has a stronger effect on the time at risk. (*Bottom*) Time at risk *t*_*r*_ is a unimodal function of “environmental tolerance” *s*_*k*_, where extinction is most likely at intermediate values. We must have 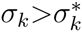 for the population to survive the initial change in the environment and 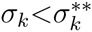 for the population abundance to drop below *N*_*c*_ (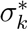 and 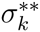 are derived by rearranging Equations 3 and 9, respectively).

### 2.2 Comparison of one-population model to previous work

Here we compare our one-population model to previous discrete-time quantitative genetic models [3, 34]. We first show how our adaptive dynamics approach gives a qualitatively similar description of trait dynamics over time and then compare our predictions of time at risk.

In a model without frequency- or density-dependence, Gomulkiewicz and Holt [3] describe the evolutionary trajectory of the population mean trait value as a geometrical approach to the optimum (Equation 5 in [3]):

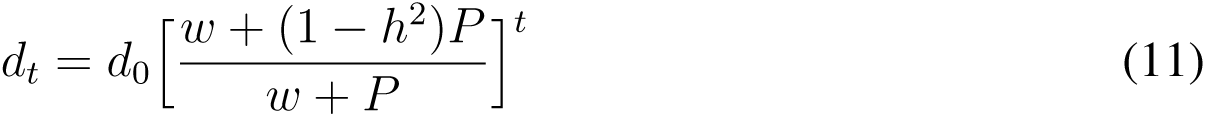

where *d*_*t*_ is the distance of the population mean trait value from the trait value giving maximum growth rate at time *t*, *w* is the variance of the growth rate function, *h*^2^ is the trait heritability, and *P* is the constant phenotypic variance [3]. We derive a qualitatively similar trajectory (Equation 7), in continuous time, from adaptive dynamics. Adaptive dynamics provides greater ecological context by including intrinsic growth rate and maximum carrying capacity as parameters in the evolutionary trajectory. The trajectories are identical when

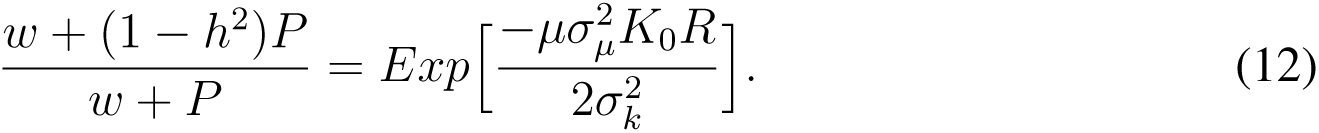

Gomulkiewicz and Holt [3] refer to Equation 12 as the evolutionary “inertia” of a trait. Inertia is bounded between zero and one in both models. When inertia is one there is no evolution. In Gomulkiewicz and Holt [3] evolution halts when trait heritability *h*^2^ or phenotypic variance is zero. In our model, inertia is determined by mutational input *µK*_0_, and evolution halts when there are no mutations. For a given *w* and *h*^2^ ≠ 0, inertia is minimized and evolution proceeds at a maximum rate in Gomulkiewicz and Holt [3] as phenotypic variance goes to infinity *P* → ∞. In our model, for a given strength of stabilizing selection per unit time 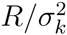,inertia to approaches zero and the rate of evolution is maximized as mutational input goes to infinity *µK*_0_ → ∞.

Note that to maintain analytical tractability both models assume the material which selection acts upon (phenotypic variance *P* or mutational input *µK*_0_) is constant. Both models will therefore be more accurate when the environmental change is relatively small. Large changes in the environment are likely to cause strong selection and large variation in abundance, which could greatly alter phenotypic variance and mutational input [30]. Since phenotypic variance and mutational input are expected to decline under strong stabilizing selection and reduced abundance [47], respectively, the analytical results of both models will tend to underestimate a population’s time at risk.

Our evolutionary trajectory aligns even closer with that of Chevin and Lande (Equation 10 in [34]; also see Equation 18a in [48]), who incorporated both density-dependence and phenotypic plasticity. The two trajectories are identical when there is constant plasticity *φ* = 0, additive genetic variance is equivalent to the supply rate of beneficial mutations times mutational size 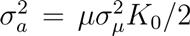 and the two measures of stabilizing selection strength per unit time are the same 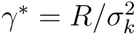.

Although our evolutionary trajectory aligns closely with those of Gomulkiewicz and Holt [3] and Chevin and Lande [34], we uncover an analytical approximation for the time at risk *t*_*r*_ by assuming a timescale separation between demographics and evolution. Gomulkiewicz and Holt [3] and Chevin and Lande [34] do not assume such a timescale separation, leading to more complex population dynamics and the need to calculate *t*_*r*_ numerically. This makes a quantitative comparison with our time at risk approximation impossible. However, Gomulkiewicz and Holt [3] agree that the time at risk *t*_*r*_ should increase with initial maladaptation (i.e., magnitude of environmental change) 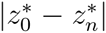 and that at high degrees of maladaptation the relationship with time at risk should be close to linear (Figure 3; Figure 5A in [3]). In addition, in both Gomulkiewicz and Holt [3] and Chevin and Lande [34] strengthening selection 1/*ω* → ∞ increases the rate of adaptation while decreasing abundance (through a decline in mean fitness). Time at risk should therefore be minimized at an intermediate selection strength, as in our model (Figure 3, bottom panel), although they do not explore this explicitly. Gomulkiewicz and Holt [3] also argue that the time at risk *t*_*r*_ should decrease with the abundance before environmental change, since the population declines geometrically beginning at this abundance. In our model, time at risk also decreases with abundance before environmental change *K*_0_, but for a different reason. Recall that because of our first-order approximation we assume a small initial stress and hence a small change in abundance. This allows us to assume that mutations are supplied at a constant rate *µK*_0_, where *µ* is the per capita mutation rate and *K*_0_ is the abundance before environmental change. A greater abundance before environmental change *K*_0_ therefore causes faster evolution resulting in less time at risk. Finally, although

### 2.3 Simulations

Adaptive dynamics assumes mutations are rare enough such that, on the timescale of evolution, the population remains monomorphic (i.e., a mutation fixes or is lost before the next arises [49]) and at demographic equilibrium (i.e., demography is faster than evolution) and that mutations are small enough to allow local stability analyses to determine evolutionary stability [40, 45]. Our approximation of time at risk *t*_*r*_ (Equation 10) also rests on the assumption that the initial stress 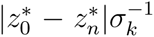 is weak. We therefore performed computer simulations to examine how well our analytical result (time at risk *t*_*r*_) holds when we relax these assumptions. To do this we varied (a) mutation rate *µ* and maximum carrying capacity *K*, (b) mutational variance 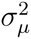, and (c) the strength of the initial stress 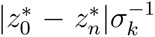 Computer simulations allow multiple phenotypes to coexist and introduces stochasticity in mutation rate and size.

Simulations describe the numerical integration of Equation 1, using a 4th-order Runge Kutta algorithm with adaptive step size, and stochastic mutations. Mutations occur in a phenotype with probability *µnDt*, where *µ* is the per capita per time mutation rate, *n* is the abundance of the phenotype, and *Dt* is the realized time step. For each mutation occuring in a phenotype with trait value *z*, one individual is given a new trait value, randomly chosen from a normal distribution with mean *z* and standard deviation *σ*_*µ*_. Trait values are rounded to the third decimal to prevent the accumulation of overly similar phenotypes. Phenotypes with abundance below one were declared extinct. Simulations began with the population at maximum carrying capacity *K* and all individuals optimally adapted with trait value 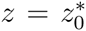 At the timestep 500, the optimal trait value instantaneously shifted to 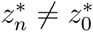. Simulations were terminated at timestep50000. Code available upon request; implemented in R [50].

Parameter values for *µ*, *K*, and 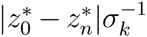 were chosen in the range of those observed for yeast exposed to increased salt concentration [5]. We estimated *σ*_*k*_ from Figure S1 in Bell and Gonzalez [5] (ESM).

In all simulations, the population evolved towards 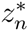 and, if successful in reaching 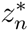 remained there. Likewise, population size always approached carrying capacity, as expected (Figure 2).

The transient dynamics, however, showed varying degrees of congruence with our predic-tion (Equations 7 and 8; Figure 4). In simulations the amount of standing phenotypic variance increases with mutation rate *µ* times population size. Our timescale assumption, which implies zero phenotypic variance, is thought to become unrealistic as *µKlog*(*K*) approaches one [51]. The threshold of *µKlog*(*K*) is obtained because *µK* is the mutational input and *log*(*K*) is the typical time of fixation for a successful mutant when the population is well adapted [51]. Over our parameter range (*µ*=*{*10^−7^, 10^−6^, 10^−5^, 10^−4^*}*, *K*=*{*10^4^, 10^5^, 10^6^*}*) *µKlog*(*K*) seemed to be an excellent predictor of accuracy; our predictions were much more accurate when *µKlog*(*K*) < 1. When *µKlog*(*K*) > 1 we greatly underestimated the time at risk (triangles in Figure 4).

**Figure 4:**
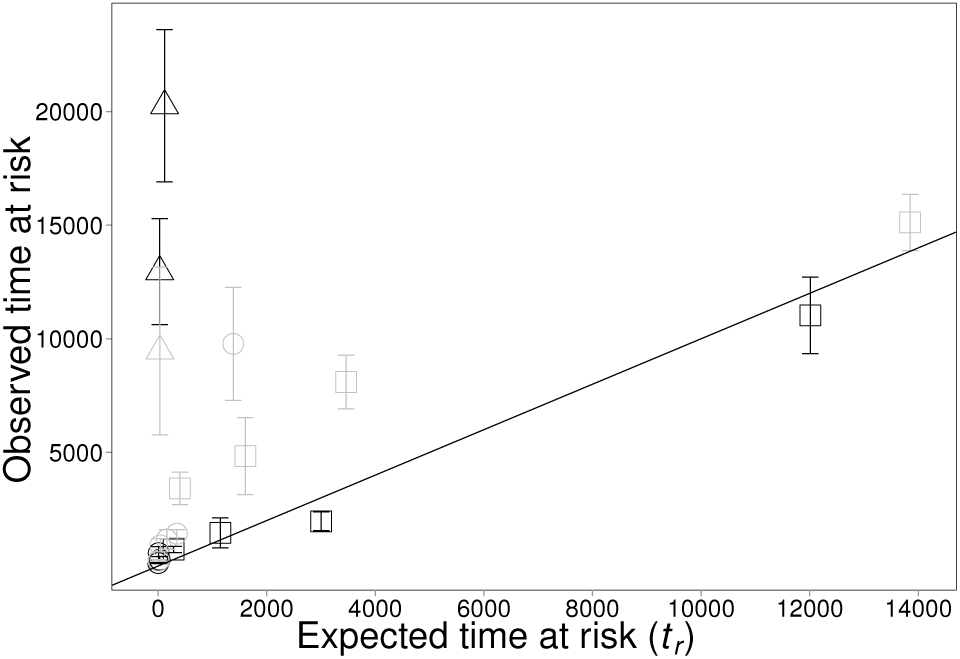
Accuracy of analytical prediction, in the one-population case. Each point represents the mean *±* SE for ten replicated simulation runs. Solid line is 1:1 line; points falling on line represent perfect predictions of time at risk *t*_*r*_. Squares: *µKlog*(*K*) *=*0.1; Circles: *µKlog*(*K*) *=*1; Triangles: *µKlog*(*K*) >1; Black: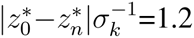 Grey: 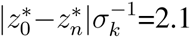 Parameters: *µ*=*{*10^−7^, 10^−6^, 10^−5^, 10^−4^*}*, *K*=*{*10^4^, 10^5^, 10^6^*}*, *s*_*µ*_=*{*0.01, 0.05*}*, *R*=1, *s*_*k*_=1, *s*_*a*_=1.5, and *N*_*c*_ is 1000 greater than the minimum abundance of each run.

Mutational variance 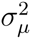 seemed to have little effect on the accuracy of our predictions, at least over the range of parameter space explored here (*σ*_*µ*_=*{*0.01, 0.05*}*; Figure 4). However, our analytical prediction did perform consistently better when the initial stress 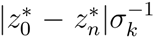 was small, for all parameter combinations (compare black 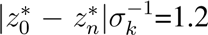 and gray 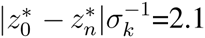 points in Figure 4).

### 2.4 Competition

We now introduce interspecific competition. Let the population dynamics of the focal population be described by the logistic growth equation:

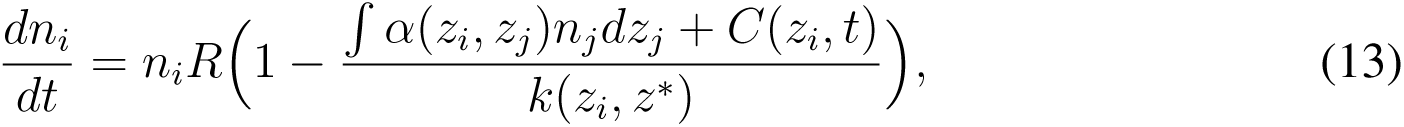

where *C*(*z*_*i*_*, t*) *=* 0 is the effect of interspecific competition on individuals in the focal population with trait value *z*_*i*_ at time *t*. We do not model the coevolution of the competitors explicitly; we instead keep interspecific competition *C*(*z*_*i*_*, t*) as general as possible, allowing it to depend on focal trait value *z*_*i*_ and vary in time *t* with any other biotic or abiotic factor (including the trait values and abundance of the focal and competing populations). For evolutionary rescue of the focal population, the only relevant dependency is with *z*_*i*_. Our formulation allows competition *C* to encompass all possible types of coevolution feedback. In fact, *C* could even be interpreted as an abiotic selection pressure. However, for brevity, we limit our discussion to *C* as the effect of a competitor. Previous studies have explicitly modeled the coevolution of competing species in a constant environment [37, 52, 53], at the expense of analytical results. All other variables in Equation 13 are defined as in the one-population case.

We again assume that mutations are rare, so that our focal population remains monomorphic with trait value 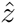 and equilibrial abundance *ñ*. In the presence of competition, equilibrium abundance of the focal population is

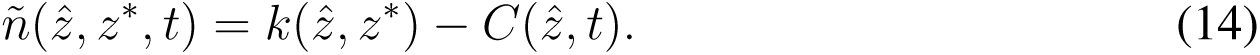

Comparison with the one-population case, where *ñ* = *K* shows how competition reduces abundance.

Now, let the competing populations coexist in a constant environment with 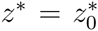 The

population will not necessarily evolve towards 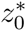 but to a “competitive optimal”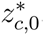 which is the trait value which maximizes equilibrial abundance *ñ* in the original environment (Appendix C). Assuming 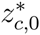 is a fitness maximum (Appendix C), the focal population will eventually evolve to the competitive optimal 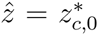. We then let the competitive optimal change abruptly, to new trait value 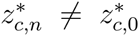. This change could arise from a shift in competion *C* or in the optimal trait value 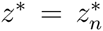 The abundance of the focal population is now 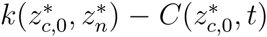 The amount of competition a population feels immediately following the environmental change 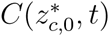 will depend on the type of environmental change as well as the response of the competitors. Competition may be close to negligible if resources remain plentiful but the abundance of competitors are greatly reduced (e.g., when a pollutant causes severe mortality in the competitor). However, competition may be exceptionally strong if the change in environment is a shift in available resources, so that the supply of resources is limiting (e.g., seed size changes on an island supporting multiple species of finch [54]). Persistence requires 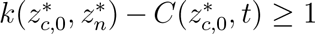1, and therefore persistence following environmental change is more likely when competition 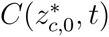 is weak.

In Appendix C we derive the local fitness gradient of the focal population. In the new environment, with 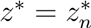 it can be written as

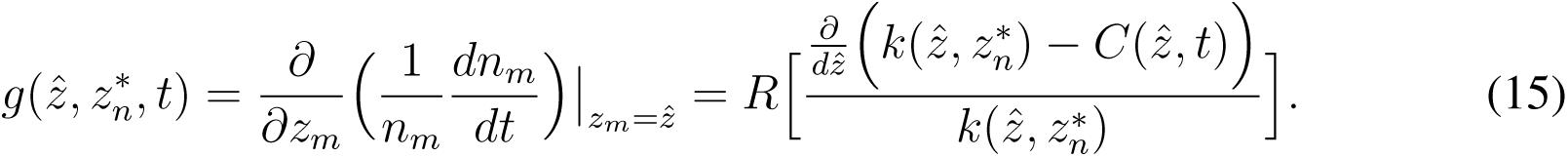

The population evolves larger population size *k - C* until 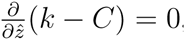 which occurs when the population reaches the competitive optimal in the new environment 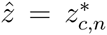 (Figure 5).

We assume that 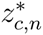 is a fitness maximum, such that the population remains monomorphic (Appendix C).

From Equation 15 we see that, relative to the one-population case (Equation 5), competition can alter the strength and direction of selection, depending on how competition changes with trait value (Figure 5). Competition increases the strength of selection when 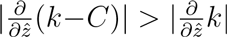 This is will always occur when competition selects in the same direction as carrying capacity (i.e., 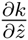 and 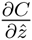 are of different signs). Competition decreases selection when 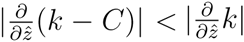 which will occur when competition weakly selects in the opposite direction to carrying capacity (i.e 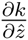 and 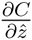 are of the same sign and 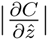 is small). When competition selects in the opposite direction as carrying capacity and has a stronger selective effect 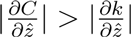 it will reverse the direction of selection and the population will evolve away from 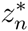 Competition has no effect on selection when it is independent of trait value 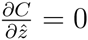

**Figure 5:**
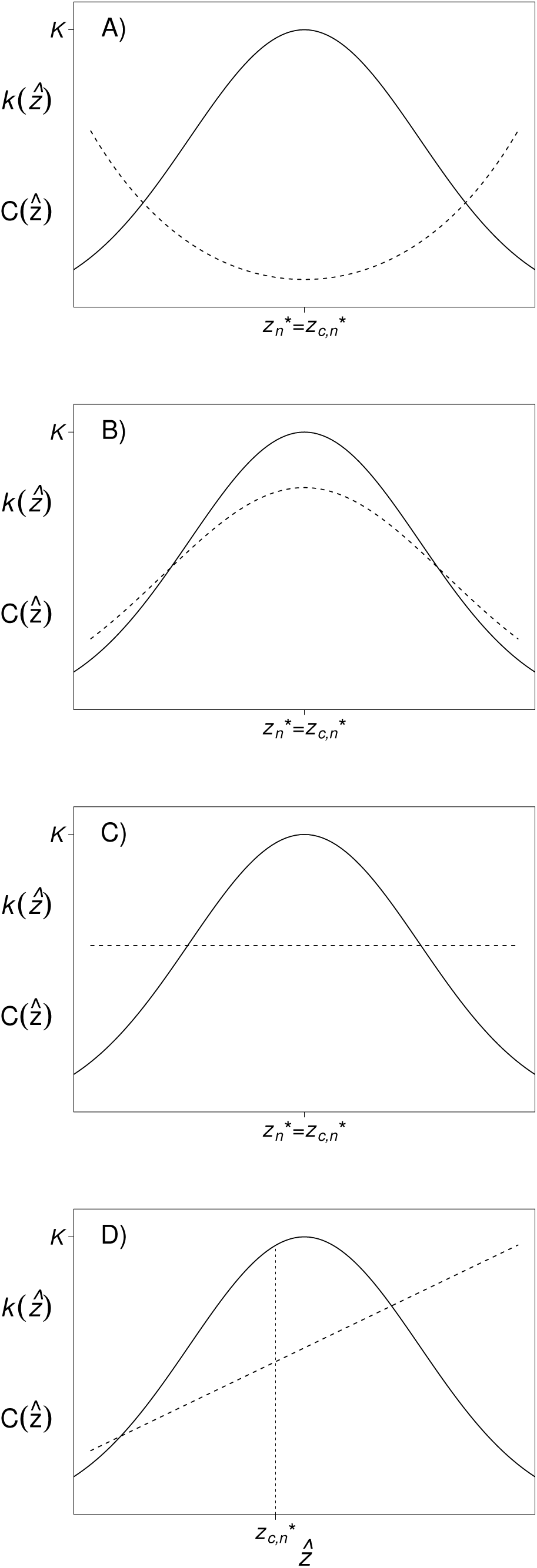
Selection pressures from carrying capacity and competition. The population evolves to increase population size according to Equation 15. Population size is carrying capacity minus competition *k - C* (solid curve minus dashed curve). Populations can persist in communities only when they have positive population size (region of persistence; solid line higher than the dashed line). The selection pressure in the new environment is proportional to the selection for carrying capacity (slope of solid curve) minus the selection for competition (slope of dashed curve). The population will therefore evolve towards the trait value for which the slopes of the two curves are equal 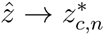. The effective selection pressure will depend on the shape of the two curves and the position of the population in trait space. (*A*) Competition increases selection pressure. Competition decreases as carrying capacity increases, meaning both carrying capacity and competition select in the same direction. (*B*) Competition reduces selection pressure. Competition increases as carrying capacity increases, meaning carrying capacity and competition exert opposing selection pressures. Note that if the competition curve was steeper than carrying capacity competition could reverse the direction of evolution. (*C*) Competition affects all phenotypes equally, and therefore has no effect on selection pressure. (*D*) Competition increases or decreases selection pressure. When 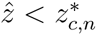 competition and carrying capacity exert opposing selection pressures. When 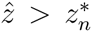 competition and carrying capacity select in the same direction, towards 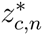 until 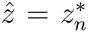 Competition and carrying capacity will then exert opposing selection pressures as the population approaches 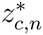

Combining Equations 14 and 15 we compute the rate of adaptation, as described by the canonical equation [44]:

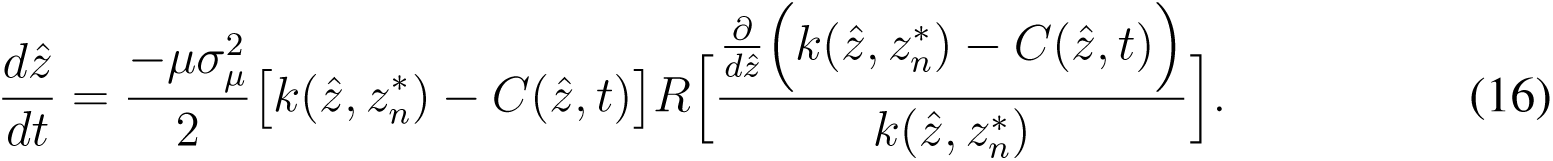

The rate the focal population adapts 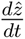 depends on how competition affects abundance relative to selection. Due to the added complexity of competition we are unable to solve Equation 16 for trait value as a function of time 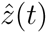 and are therefore unable to compute a time at risk *t*_*r*_, as we did in the one-population case. However, we can show when competition will help or hinder adaptation, and therefore when competition has the potential to increase or decrease the likelihood of evolutionary rescue. Rearranging Equation 16 and comparing to the one-population case (Equation 6) shows that competition will increase the rate of adaptation when (Appendix D)

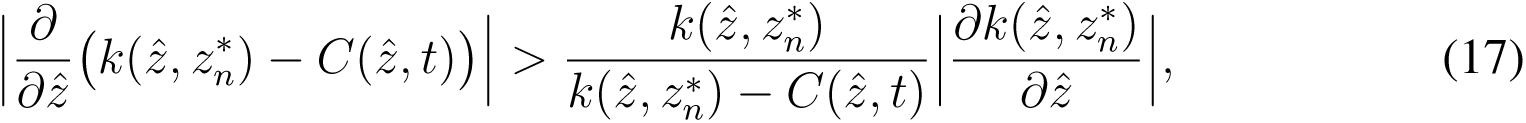

and decrease the rate of adaptation when the inequality is reversed. Competition will tend to speed adaptation when competition *C* is weak and gets much weaker as the focal population evolves towards 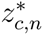(dot-dashed curve in Figure 6). Note that although competition may increase the rate of adaptation, and therefore cause a greater *rate of increase* in abundance, abundance will still be depressed by competition. Competition’s effect on evolutionary rescue (the time at risk *t*_*r*_) will therefore depend on both its effect on adaptation and the abundance *k − C* relative to critical abundance *N*_*c*_ (bottom panel in Figure 6). As maximal abundance *K − C* approaches the critical value *N*_*c*_ evolutionary rescue becomes less likely, and regardless of the rate of adaptation, when *K − C ≤ N*_*c*_ evolutionary rescue is impossible.

**Figure 6:**
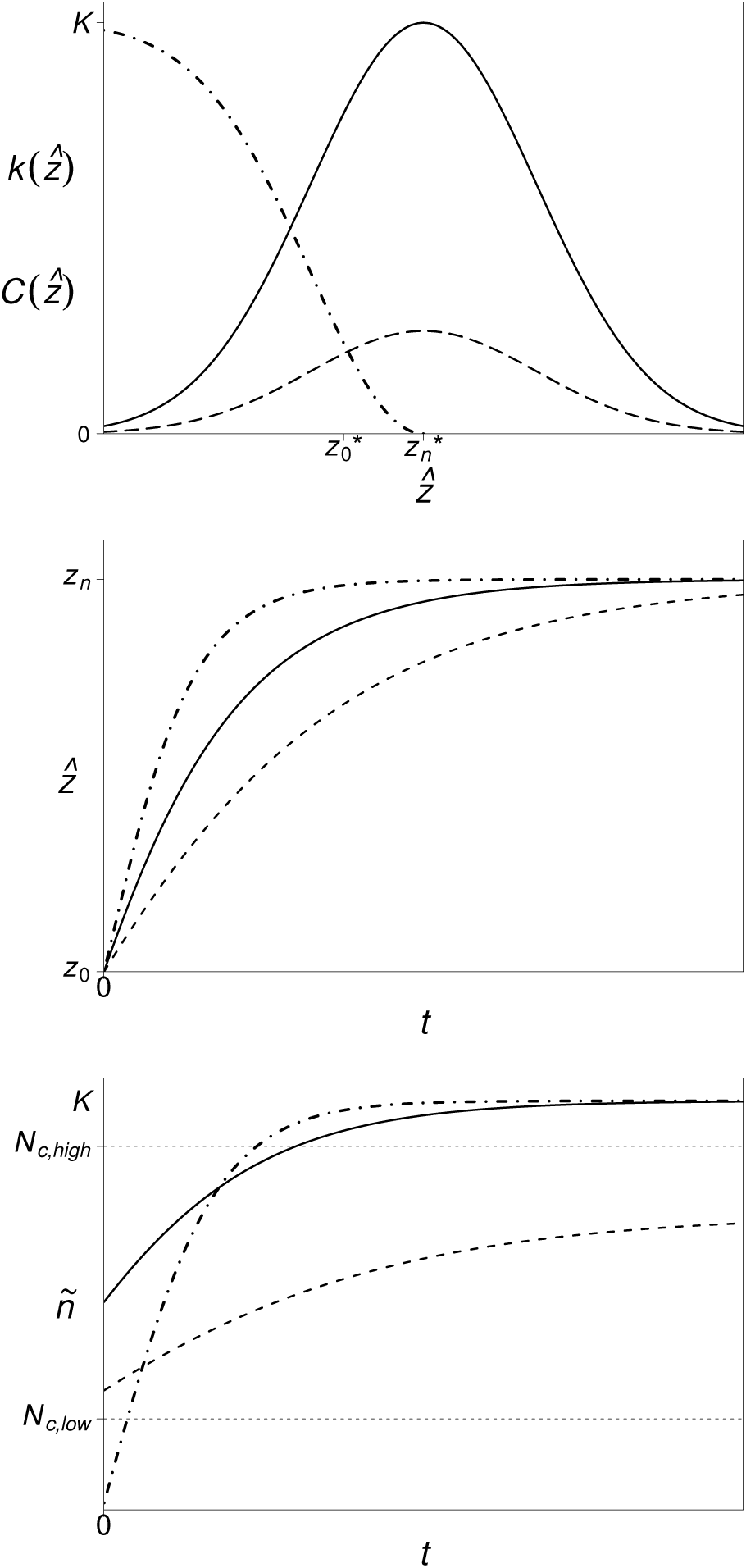
Competition can help or hinder evolutionary rescue. (*Top*) Carrying capacity *k* (solid curve) as a function of trait value 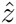 and two competition *C* scenarios: complete niche overlap (dashed curve) or partial niche overlap (dot-dashed curve). (*Middle*) With complete niche overlap (dashed curve) competition increases as the population adapts, and the population therefore adapts slower than it would without competition (solid curve). With partial niche overlap (dot-dashed curve) competition decreases as the population adapts, and the population therefore adapts faster. (*Bottom*) The time a population spends at risk of extinction (the time abundance *ñ* is below critical abundance *N*_*c*_) depends on competition’s effect on abundance and evolution as well as on the value of the critical abundance. For instance, when the critical abundance is low *N*_*c,low*_ both competition scenarios increase the time at risk relative to when there is no competition (solid curve) because they depress the focal population’s abundance. However, when the critical abundance is high *N*_*c,high*_ partial niche overlap (dot-dashed curve) decreases the time at risk relative to the no competition case (solid curve) because it sufficiently increases the rate of adaptation.

INCLUDE FIGURE 6 HERE

## 3 Discussion

In nature, population abundance cannot increase indefinitely [55]. One of the main “checks of increase” [56] is competition for resources [17, 57, 19, 58, 59]. Because populations with lower abundances are more likely to go extinct [46], any factor which limits abundance is likely to hinder persistence, especially when the environment changes [22]. However, when we consider that populations can persist in new environments by adapting [3, 5], competition has a second effect, in addition to lowering population size, which could potentially help populations persist in novel environments. Since the rate a population adapts depends on the strength of selection it experiences [60, 44], competition which increases the strength of selection may speed-up adaptation [61] possibly increasing the chances of persistence in the face of change.

Intraspecific competition often has relatively little impact on selective pressures [58, 62] (but see [63]) and therefore the effect it has on evolutionary rescue will often be determined primarily by the effect it has on abundance. Previous computer simulations have suggested that negative density-dependence will have little effect on population persistence because survival depends on the dynamics of populations which are well below carrying capacity [29]. More recent analytical work has come to a different conclusion, showing that, relative to the density-independent case, density-dependence can increase the rate at which abundance declines as well as decrease the rate abundance recovers, therefore increasing the time a population spends at risk of extinction [34]. The conflicting results are due to the different types of density-dependence used in the two studies. In Boulding and Hay [29] density-dependence is linear (i.e., per capita growth rate declines linearly with abundance) while in Chevin and Lande [34] density-dependence is stronger than linear at low abundances (the per capita growth rate declines logarithmically with abundance). Since it is the effect of density-dependence at low abundances that is critical for population persistence, this explains why Chevin and Lande [34] claim density-dependence increases the chances of extinction. A similar trend is expected in biological invasions, where populations experiencing strong density-dependence at low abundances are predicted to invade slowly [64].

Here we assume evolution is slow, and hence, on the timescale of evolution, populations are always at carrying capacity. Carrying capacity therefore indicates how well a population is adapted; populations below carrying capacity will increase in abundance without evolving, and hence may not require evolutionary rescue if their carrying capacity is large enough. In our model, it is the *maximum* carrying capacity that affects the potential, and need, for evolutionary rescue. Since abundance asymptotically approaches maximum carrying capacity in evolutionary time (Figure 2), maximum carrying capacity will have a larger effect on the time at risk as it approaches the critical abundance (Figure 3).

Notice that maximum carrying capacity plays both a demographic and evolutionary role; for a given environmental change, larger values keep populations at larger abundances (*K* in Equation 8) and, following the change, increase the rate of evolution (*K*_0_ in Equation 7). Here we assume greater abundances lead to faster evolution because they cause greater mutational inputs. In previous models (e.g., [3, 34]), where the rate of evolution is determined by additive genetic variation instead of mutational input, the relationship between population size and the rate of evolution can be weaker (reviewed in [65]). Although non-additive genetic effects, such as epistasis and dominance, and temporal fluctuations in abundance (leading to lower effective population sizes) can weaken the relationship between population size and the rate of evolution [66], they do not qualitatively alter our results, but merely lead to a slower rate of evolution than predicted.

Given the differences between quantitative genetics and adaptive dynamics [51], our results are surprisingly consistent with previous quantitative genetic models of evolutionary rescue (e.g., [3, 34]). We derive a similar evolutionary trajectory and agree with Gomulkiewicz and Holt [3] on with how time at risk should increase with initial maladaptation and decrease with abundance before environmental change.

There is, however, one major difference between our approach and previous models of evolutionary rescue. All previous models assume the environmental change affects intrinsic growth rate, and that it is the intrinsic growth rate that must evolve fast enough to allow persistence. In our model, intrinsic growth rate *R* has no effect on abundance since populations are assumed to remain at demographic equilibrium, which is independent of *R*. In particular, the environmental change might affect *R* with no effect on abundance (so long as *R >* 0). Intrinsic growth rate is therefore irrelevant for evolutionary rescue in our model. Here rescue depends on the effect of the environmental change on carrying capacity *k*, and the evolution of *k*. Past models describe evolutionary rescue under *r*-selection while we describe evolutionary rescue under *K*-selection [67, 68]. Hence, our model is more applicable to situations where density-dependence remains strong following the environmental change, during subsequent adaptation. Density-dependence will remain strong when the demand for resources continues to equals the supply. Obviously, density-dependence will remain strong when an environmental change acts only to reduce the supply of resources. This describes how a population of Darwin’s finches has responded to drought [54]. The drought lowered the supply of seeds the finches ate, causing a rapid decline in finch abundance. Competition for small seeds intensified following drought and the finch population remained at carrying capacity, a carrying capacity which had been reduced by decreased food supply. Density-dependence can also be maintained when an environmental change leaves the supply of resources unaffected but increases the per capita demands. For instance, if stress tolerance requires increased energetic demands, a population exposed to a stress may continue to experience strong density-dependence despite a decline in abundance and unaffected resources. This may describe the situation observed in recent experiments of evolutionary rescue in yeast populations exposed to salt, where glucose concentration was unaffected [5, 38].

Simulations indicate that our analytical approximations are sensitive to mutational input and the fixation times of new beneficial mutations. When mutations are too frequent or fixation times are too long we consistently underestimate the time at risk (Figure 4). The underestimate likely arises from the adaptive dynamic assumption that fixation occurs instantaneously and the population remains monomorphic. In simulations which permit greater polymorphism, less fit phenotypes compete with those closer to the adaptive optimum, imposing a demographic load on the population. The continued existence of less fit phenotypes slows the increase of carrying capacity, causing populations to remain at risk of extinction for longer than expected. This is similar to what, in microbial evolution, is refered to as “clonal interference” [69]. However, many populations should conform to our low mutation input assumption. For instance, the mutations rate of *Saccharomyces cerevisiae* salt tolerance is approximately *µ* = 10^−7^ mutations per genome per generation [5]. Since our analytical approximations are accurate when *µKlog*(*K*) < 1, our method can handle yeast populations of about one million cells or less.

Although our approximations are most sensitive to high mutational inputs and slow fixation times, our assumption that mutational input is constant throughout adaptation (similar to assuming constant phenotypic variance [48, 3]) becomes less realistic as the initial stress becomes larger (Figure 4). Assuming constant mutational input is necessary for an analytical solution, but causes us to consistently underestimate the time at risk. In reality, environmental changes will cause reductions in abundance which will decrease the supply rate of new mutations (or phenotypic variance [48]), effectively “pulling the rug out from under evolutionary rescue” [30]. Both ours and the traditional quantitative genetic [48] analytical approximations are less accurate under strong selection [29]. Because high mutation rates, long fixation times, and large initial stresses all cause our approximation to underestimate the time at risk, our analytical results can be considered a best-case scenario for population persistence.

Competition between individuals of distinct species is likely to cause dramatic changes in selective pressures [70, 62]. If competition is strong enough to drive rapid adaptation, competitors can potentially help a population adapt and persist following an environmental change. In a continuously changing environment, computer simulations of two competing populations have shown that competition can aid the persistence of the better adapted population by increasing selective pressure, effectively “pushing” the phenotype of the better adapted population toward the moving optimal [36]. Our results clarify this point - competition can aid population persistence when it increases the selective pressure to evolve to the new environment - and give an additional requirement: competition must increase selection pressure enough to overcome the negative effect of reduced abundance. The effect of competition on evolutionary rescue can be explained in terms of the overlap between the competitor’s niche and the niche the focal population is attempting to adapt to. When the focal population is forced to adapt to a niche already occupied by a competitor (strong niche overlap), competition will hinder adaptation because competition selects in the opposite direction as the new environment (dashed curve in Figure 6). On the other hand, when the competitor has a niche which only partially overlaps the niche the focal population is attempting to adapt to, it can speed adaptation by depressing the fitness of individuals in the focal population which are farther from the new niche (dot-dashed curve in Figure 6). We can illustrate this concept by returning to the example of Darwin’s finches. Drought reduced the supply of small seeds, shifting the niche available to the medium ground finch (*Geospiza fortis*) to larger seeds. In general, this caused *fortis* populations to evolve to larger size [54]. However, in the presence of the large ground finch *G. magnirostris*, who eat large seeds (strong niche overlap), larger *fortis* were outcompeted by *magnirostris*, preventing *fortis* from evolving to larger size [71, 72]. Meanwhile, in the presence of the small ground finch *G. fuliginosa*, who eat small seeds (partial niche overlap), smaller *fortis* were outcompeted by *fuliginosa*, causing *fortis* to evolve to a larger size faster than they did in the absence of competitors [61]. Populations of *fortis* approached the new adaptive peak faster when in competition with *fuliginosa* because *fuliginosa* increased selection pressure towards the peak. What remains to be seen, and what is pivotal for evolutionary rescue, is whether the increased adaptation of *fortis* in the presence of *fuliginosa* overcame the reduction in *fortis* abundance caused by competition with *fuliginosa*.

On the other hand, competition may be the very reason evolutionary rescue is required for persistence in the first place. Invasive species, for example, can greatly reduce the abundance of pre-existing competitors, putting many populations at risk of extinction (reviewed in [14]). Our results suggest that some invading populations, which are themselves the cause of extinction risk, hinder evolutionary rescue in their competitors, while other invaders may permit rapid adaptation. The model presented here may therefore help predict if an invasive species is likely to cause niche displacement or extinction (reviewed in [13]). Since few examples of extinction are associated with competitive interactions between native and invasive species [13], invading competitors may often allow rapid adaptation.

Although we have shown that competition can help evolutionary rescue under specific circumstances, we have simultaneously shown that in other circumstances competition will surely hinder persistence. Interspecific competition is also expected to reduce rates of adaptation in the context of species’ range limits [72] and gradual environmental changes in metacommunities [73]. When competition hinders adaptation, we expect evolutionary rescue to be more common in communities with reduced niche overlap, [74] or greater character displacement [75], since in these communities there should be less interspecific competition.

Coevolution can alter the demographic costs and selection pressures imposed by competition, therefore impacting population persistence [70]. In our case, altering the strength and selection pressure of competition means a shift in the height and slope of the competition curve (Figure 5), respectively, as the focal population evolves. A number of previous studies have investigated the effect of coevolution between competitors (although not in the context of evolutionary rescue; [37, 52, 53]). Here, instead of asking how a specific form of coevolution influences persistence, we ask a more general question: what types of coevolution help (or hinder) evolutionary rescue? For example, if coevolution is expected to cause strong character displacement [53], not only will the less adapted population “push” the better adapted population to even greater levels of adaptation, but the better adapted population will also “push” the less adapted population away from it, reducing the positive effect of competition on evolutionary rescue.

Although our analytical approach sometimes requires stricter assumptions than simulation studies (e.g., constant mutational input), it avoids the finite choice of parameter values demanded in simulation studies, and thereby provides more general results. For instance, our expression for time at risk (Equation 10) shows a unimodal relationship with environmental tolerance (Figure 5), indicating that extinction is most likely at intermediate tolerances. Extinction is most probable at intermediate environmental tolerances because small tolerances cause strong selection pressures and hence - if the population can survive the initial stress - fast evolution, while large tolerances allow high degrees of maladaptation without a demographic cost. To our knowledge, this is the first time this relationship has been clearly demonstrated.

In a recent experiment of adaptation to a novel environment under competition, Collins [9] subjected pairs of competing photosynthetic microbe strains to increased carbon dioxide levels. Despite the loss of one of the competing strains part way through the experiment, the presence of a competitor at the beginning of the experiment always reduced the final abundance of the survivor. Collins [9] partitioned the effects of physiology, evolution to increased carbon dioxide levels, and competitive ability on final abundance. She found that when competition had an effect it was always opposing evolution to carbon dioxide. In other words, when competition affected adaptation it was because the superior competitor went extinct while the strain most capable of adapting to the new environment evolved slower than it would have in monoculture. A trade-off between competitive ability and the ability to adapt to abiotic change lowered the abundance of both strains, impeding evolutionary rescue of all. In our model, this amounts to a positive correlation between carrying capacity and competition during the initial stages of adaptation. When this positive correlation exists, competition will nearly always impede evolutionary rescue.

To our knowledge, this is the first analytical work to investigate the effect of interspecific competition on evolutionary rescue following an abrupt environmental change. In doing so, we have highlighted the general ecological and evolutionary settings where competition should help or hinder persistence to environmental change.

## 4 Acknowledgements

We thank Helene Weigang, Ophélie Ronce, Peter Jackson, Robert D. Holt, and an anony-mous reviewer for helpful comments on the manuscript. MMO was funded by a Alexander Graham Bell Canada Graduate Scholarship from the National Sciences and Engineering Research Council of Canada, the Quebec Centre for Biodiversity Science, and the Dr. Neal Simon Memorial Scholarship. CdM acknowledges a Discovery Grant from the Natural Sciences and Engineering Research Council of Canada.

## 5 Appendix A

Here we find the singular strategy in the one-population case and evaluate its stability. Detailed methods can be found in Geritz et al. [40]. From Equation 1 the local fitness gradient is

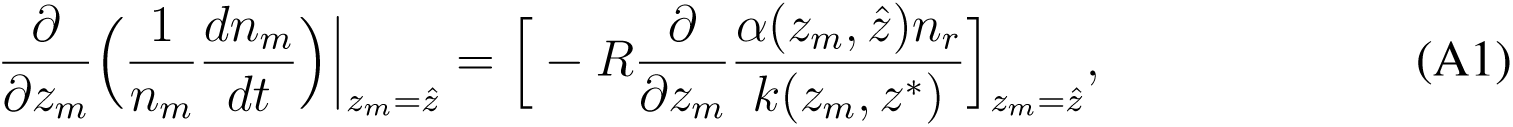

where *z*_*m*_ is the trait value of a rare mutant with abundance *n*_*m*_ and 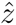 is the trait value of the resident with abundance *n*_*r*_. Dropping the arguments of the functions and denoting 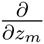 with prime gives

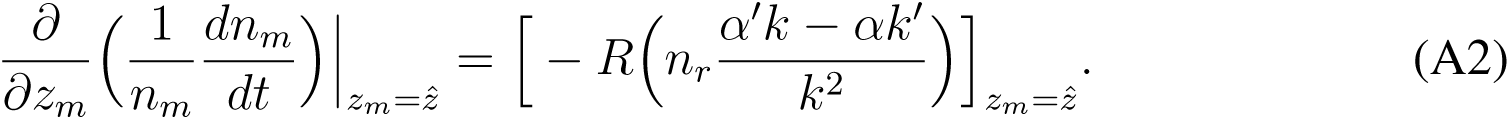

Assuming 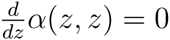 and *α*(*z, z*) = 1, evaluating at 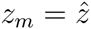 gives

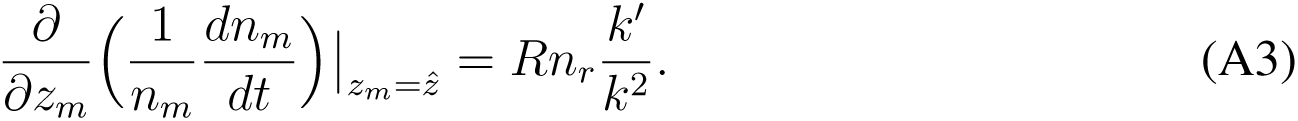

Specifying *k* as a Gaussian function (Equation 2) with mean *z*^*\**^ and variance 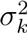,

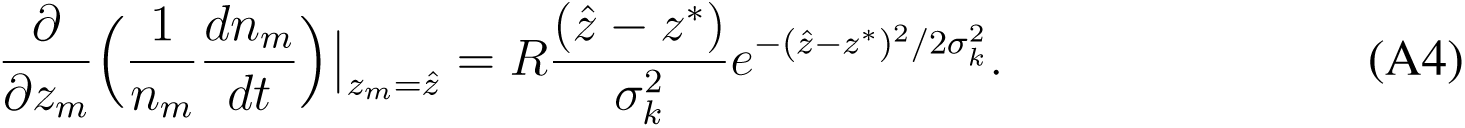

The local fitness gradient is zero when 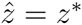 (i.e., *z*^*\**^ is the singular strategy). If *z*^*\**^ maximizes the local fitness gradient it is a fitness maximum and therefore evolutionary stable (ESS). If *z*^*\**^ minimizes the local fitness gradient it is a fitness minima and evolutionary branching may occur [40]. The singular strategy is a fitness maximum when

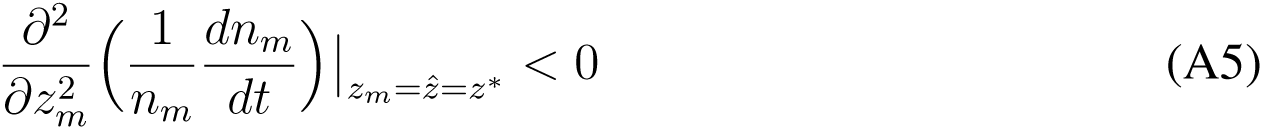

or, equivalently

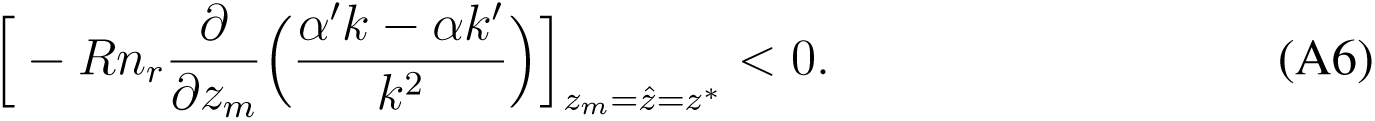

Evaluating at 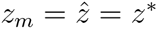 gives

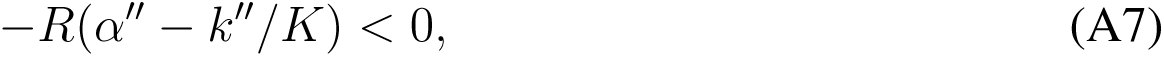

and *z*^*\**^ is therefore evolutionary stable when

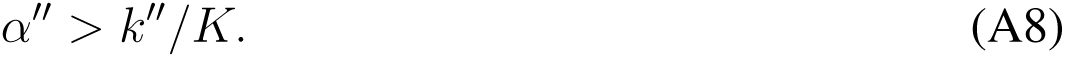

Specifying *k* as Equation 2, *z*^*\**^ is evolutionary stable when

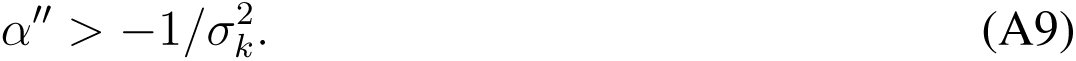

The population will converge on the singular strategy *z*^*\**^ only if

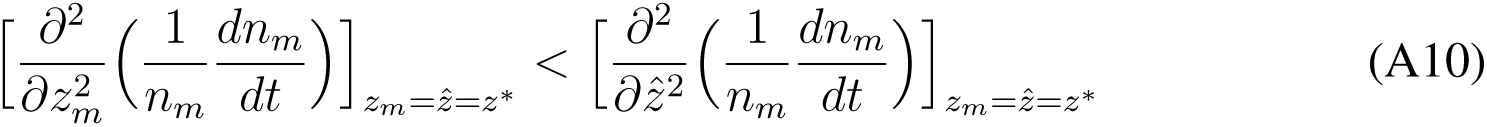

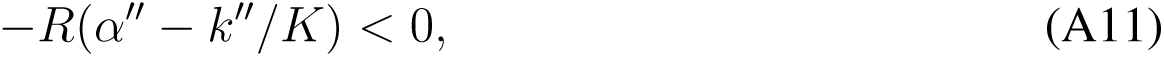

and so, if the singular point is evolutionary stable it is also convergence stable. Throughout the paper we assume Equation A11 holds to simplify our analysis of evolutionary rescue.

## 6 Appendix B

Here we derive approximations for the ecological and evolutionary dynamics in the one-population case (Equations 7 and 8). We first move all terms of Equation 6 with 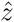 to the left-hand side and bring *dt* to the right. Then taking the integral,

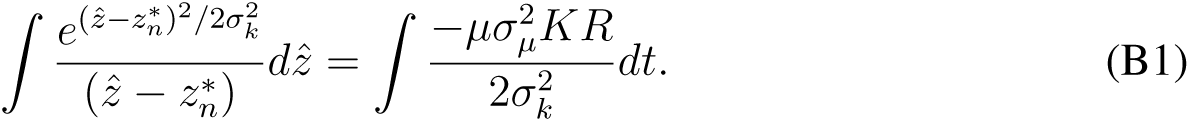

Since there is no analytical solution for the indefinite integral on the left hand side, we use the Taylor expansion about 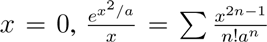 with 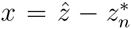 and 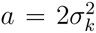 Taking the Taylor series about 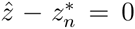 leads us to assume a small change in abundance and hence constant mutational input *µK*. We therefore replace *K* with *K*_0_ to indicate that mutational input depends on the original abundance. We now have

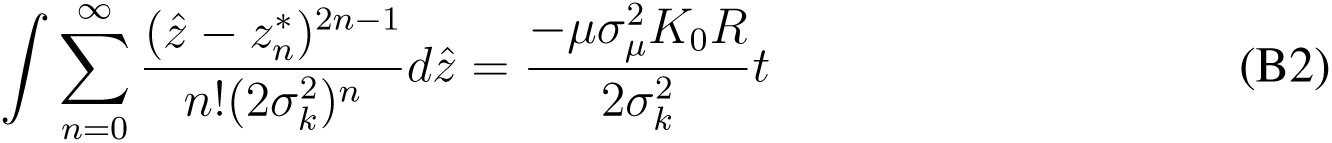

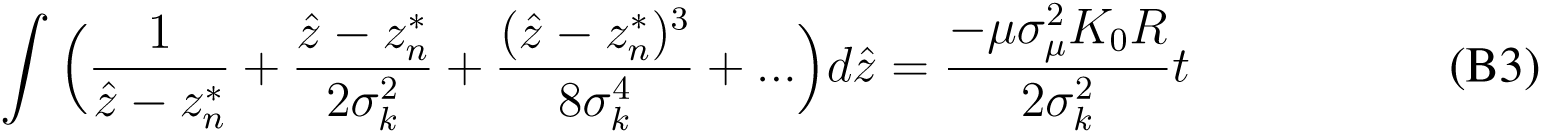

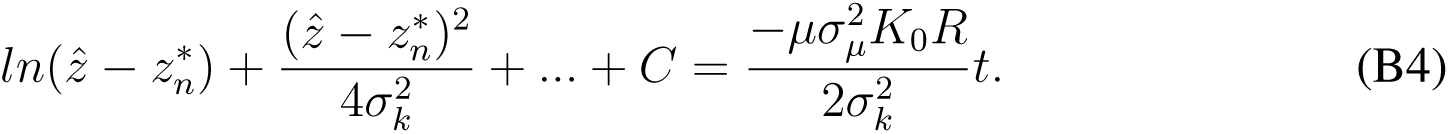

Approximating to the first order

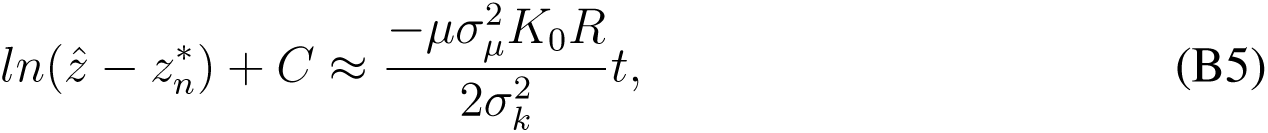

and solving for 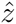 gives

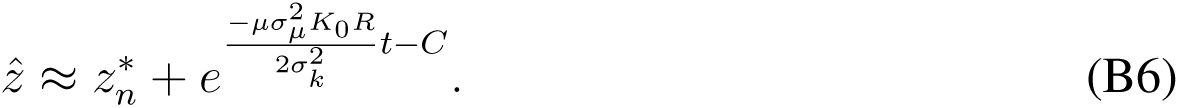

At *t* = 0 we have 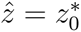 so 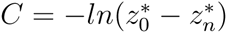 and we get Equation 7:

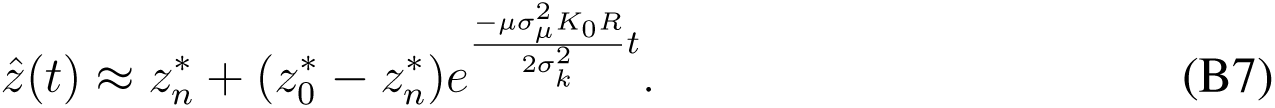

Subbing Equation B7 into Equation 2 gives an approximate description of population abundance across evolutionary time (Equation 8).

## 7 Appendix C

Here we find the singular strategies for a population experiencing interspecific competition and evaluate their stability. From Equation 13 the local fitness gradient is

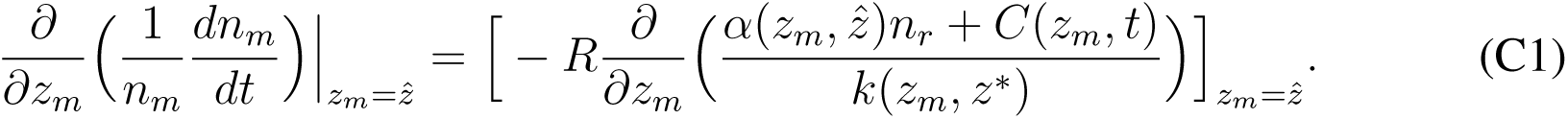

where *z*_*m*_ and *n*_*m*_ are the trait value and abundance of a rare mutant, respectively, in a population with resident trait value 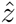 and abundance *n*_*r*_. We drop the arguments of the functions and denote 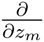 with prime. Expanding gives

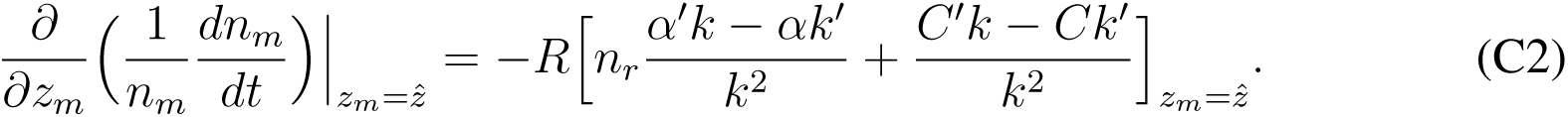

And from Equation 14:

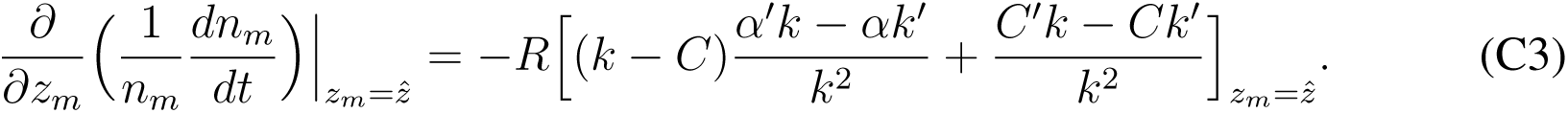

Evaluating at 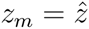:

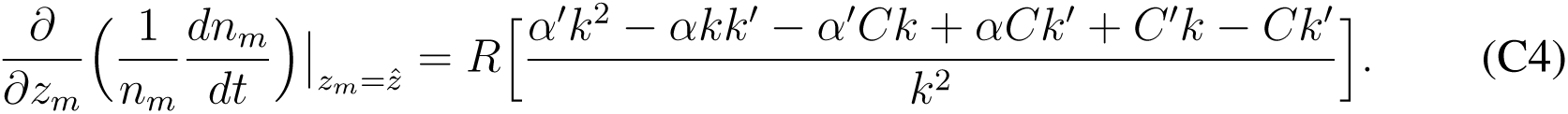

Assuming intraspecific competition *α* is maximal when individuals share the same trait value,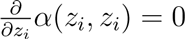 and _(zi; zi) = 1

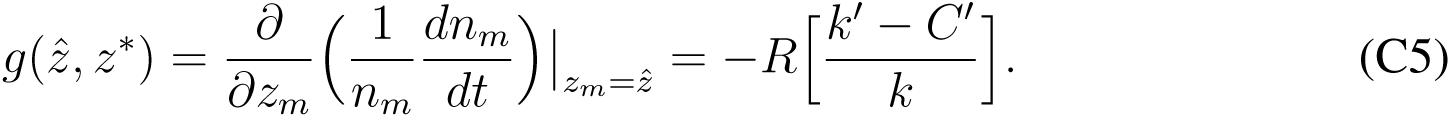

Equation C5 determines the direction of selection. Evolution proceeds until 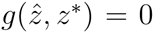, in this case when *k*′ = *C*′. The trait values giving 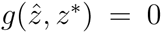 are evolutionary singular strategies, which we will denote 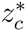 If 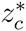 maximizes 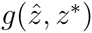, 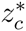 is a fitness maximum; when 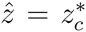 no nearby mutant can invade and the population remains monomorphic with 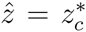 However, when 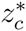 minimizes 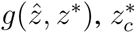 is a fitness minima and evolutionary branching may occur [40]. A singular point 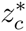 is a fitness maximum when

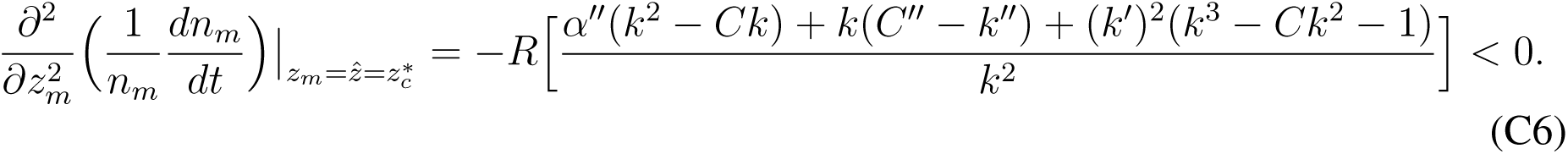

To simplify our analysis of evolutionary rescue we assume that all singular strategies our population approaches are fitness maxima. This assumes, at 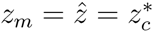

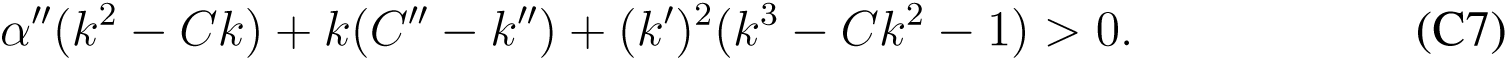

We will also assume the singular strategies are convergence stable, requiring:

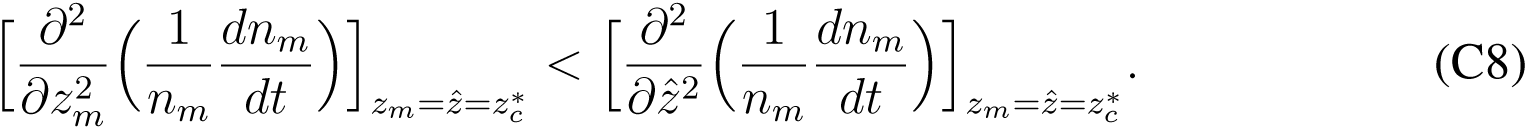

## 8 Appendix D

Beginning with Equation 16, we look to find when interspecific competition speeds adaptation towards the optimal 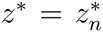 Dropping the arguments of the functions and denoting 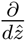 with prime, Equation 16 reads

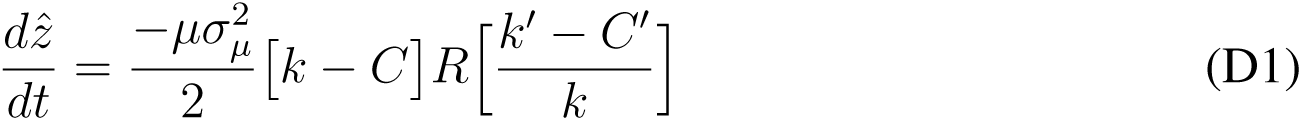

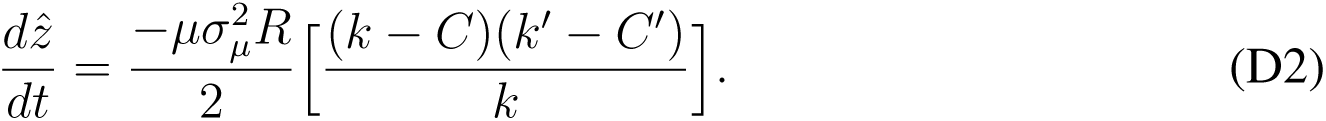

Since in the one-population case 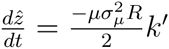 (Equation 6), competition will speed evolution

When

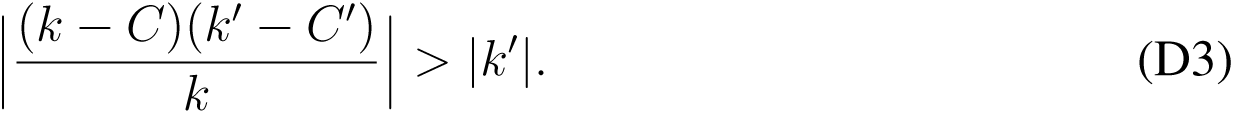

Since *k* and *k - C* must be positive for the population to persist,

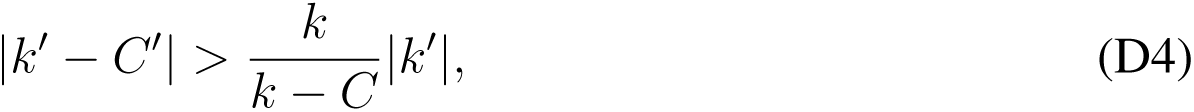

yielding Equation 17.

